# Predicting 3D genome folding from DNA sequence

**DOI:** 10.1101/800060

**Authors:** Geoff Fudenberg, David R. Kelley, Katherine S. Pollard

## Abstract

In interphase, the human genome sequence folds in three dimensions into a rich variety of locus-specific contact patterns. Here we present a deep convolutional neural network, Akita, that accurately predicts genome folding from DNA sequence alone. Representations learned by Akita underscore the importance of CTCF and reveal a complex grammar underlying genome folding. Akita enables rapid *in silico* predictions for sequence mutagenesis, genome folding across species, and genetic variants.

## Main text

Recent research has advanced our understanding of the proteins driving and the sequences underpinning 3D genome folding in mammalian interphase, including the interplay between CTCF and cohesin^1^, and their roles in development and disease^2^. Still, while disruptions of single bases can alter genome folding, in other cases genome folding is surprisingly resilient to large-scale deletions and structural variants^3,4^. As follows, predicting the consequences of perturbing any individual CTCF site, or other regulatory element, on local genome folding remains a challenge.

Previous machine learning approaches have either: (1) relied on epigenomic information as inputs^5–7^, which does not readily allow for predicting effects of DNA variants, or (2) predicted derived features of genome folding (e.g. peaks^8,9^), which depend heavily on minor algorithmic differences^10^. Making quantitative predictions from sequence poses a substantial challenge: base pair information must be propagated to megabase scales where locus-specific patterns become salient in chromosome contact maps.

Convolutional neural networks (CNNs) have emerged as powerful tools for modelling genomic data as a function of DNA sequence, directly learning DNA sequence features from the data. CNNs now make state-of-the-art predictions for transcription factor binding, DNA accessibility, transcription, and RNA-binding^11–14^. DNA sequence features learned by CNNs can be subsequently post-processed into interpretable forms^15^. Recently, Basenji^16^ demonstrated that CNNs can process very long sequences (∼131kb) to learn distal regulatory element influences, suggesting that genome folding could be tractable with CNNs.

Here we present Akita, a deep CNN to transform input DNA sequence into predicted locus-specific genome folding. Akita takes in ∼1Mb (2^20^ bp) of DNA sequence and predicts contact frequency maps for all pairs of ∼2kb (2048bp) bins within this region. Crucially, this allows Akita to predict the effects of mutating single base pairs. We trained Akita with five of the highest-quality Hi-C and Micro-C datasets as targets (**Table 1**), focusing on the locus-specific patterns evident in log(observed/expected) maps, minimizing the mean squared error (MSE) between predictions and targets.

The Akita architecture consists of a ‘trunk’ based on the Basenji^16,17^ architecture to obtain 1D representations of genomic sequence, followed by a ‘head’ to transform to 2D maps of genome folding (**Fig. 1a, Methods**). In the ‘head’, we first averaged the representations of genomic bins *i* and *j*. Averaging produced slightly better generalization accuracy relative to several alternatives, including concatenation (**Supplemental Fig. 1, Supplemental Note**). As genomic distance can impact regulatory element communication, we appended a positional encoding of the distance between bins. Drawing inspiration from CNNs used in image processing, we computed multiple layers of dilated residual 2D convolutions, re-symmetrizing after each block. Finally, we compared the upper triangular regions of target and predicted maps. We reasoned the trunk would enable Akita to learn DNA motifs and how they combine into a grammar for genome folding. In turn, the head would recognize relationships between these features and propagate this information across the map, while accounting for the dependencies between neighboring bins.

**Figure 1:**
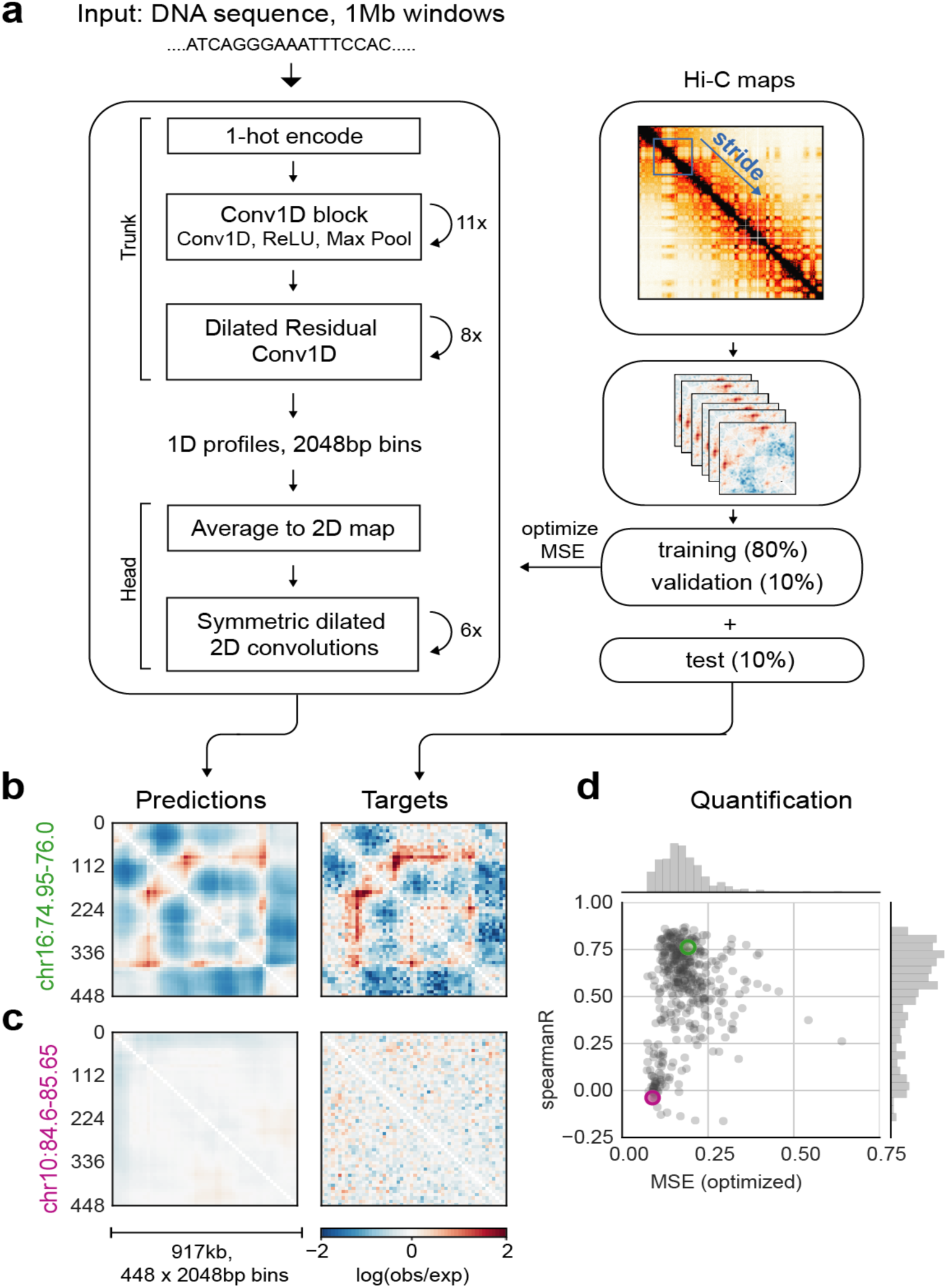
Akita makes locus-specific predictions for 3D genome folding from DNA sequence. **a**. Akita consists of a ‘trunk,’ based on the Basenji architecture^16^, followed by a ‘head’ to transform to 2D maps of genome folding. The trunk involves: (i) input 1Mb of 1-hot encoded DNA; (ii) 1D convolution trunk, where each block performs a max pool operation between adjacent positions to iteratively reduce to a bin size of 2048 bp (iii) dilated residual 1D convolutions to propagate local information across the sequence. The ‘head’ involves: (i) forming 2D maps from the 1D vectors by averaging each pair of vectors at positions (*i, j*); (ii) symmetric dilated residual 2D convolutions; (iii) dense layer with linear activation to predict log(observed/expected) chromosome contact maps. We considered 2048bp binned maps, as high-quality Hi-C and Micro-C datasets ascertain genome folding at this resolution with tractable technical variance. We compared upper triangular regions of maps cropped by 32 bins on each side, making symmetric predictions for 448×448 bin (∼917kb) maps. We trained our model on regions of the genome obtained by striding along Hi-C maps, using an 80/10/10 training/validation/test split. **b,c**. Predicted and experimental log(observed/expected) contact frequency for two representative regions in the test set for Human Foreskin Fibroblast (HFF) Micro-C ^20^. **d**. Quantification for the held-out test set: mean-squared error (MSE), which we optimize in model training, versus Spearman R, both calculated per region for each pair of targets and predictions for HFF Micro-C. Green and purple circles show regions from (b) and (c). Note correlations display a bimodal shape: regions with few locus-specific features have low MSE and low Spearman R.

Akita learned a predictive representation of genome folding from DNA sequence (overall 0.14 MSE, 0.61 Pearson, 0.56 Spearman on held-out test data). On a region-by-region basis, Akita captured the variety of patterns seen experimentally (**Fig. 1b,c**) and displayed a bimodal distribution of correlations. Many of the lower correlations represented correct predictions for featureless experimental maps, indicating that correlations, while interpretable, underestimated model performance on this task. Indeed, Akita’s predictions also captured the strength of locus-specific folding seen experimentally (**Supplemental Fig. 2**). By simultaneously training on all five datasets in a multi-task framework, Akita has greater accuracy for each dataset compared to models trained on that dataset alone (**Supplemental Fig. 3**). Still, Akita predicted limited cell-type-specific differences (**Supplemental Fig. 4**). We hypothesize this was constrained by the extent of cell-type specific differences currently ascertainable in experiments. Even the dramatic cellular transformation of cardiomyocyte differentiation displayed minimal differences upon exit from the ESC state and mainly similarities thereafter via Hi-C^18,19^. Unless noted, we thus focused our following analyses on Akita’s predicted outputs for HFF Micro-C^20^, the training dataset with the strongest locus-specific folding.

Akita predicted more prominent patterns in regions with greater CTCF binding and DNAse hypersensitivity (**Supplemental Fig. 2**). Visually, salient patterns in predicted maps often aligned with CTCF ChIP-seq peaks (**Fig. 2b**). However, CTCF motifs were too prevalent to observe a correspondence at the bin level (**Supplemental Fig. 5**). Fortunately, Akita enabled us to ascertain their influence via *in silico* mutagenesis; while training Akita was computationally intensive, effects of sequence changes can be predicted in seconds. Akita predicted greatly diminished locus-specific patterns upon CTCF motif mutagenesis (**Fig. 2e**). Still, Akita predicted some patterns would persist, and these often aligned with DNase hypersensitive sites that lacked evidence of strong CTCF binding. Inverting all CTCF motifs produced very different predictions, redistributing rather than abrogating contact patterns (**Fig. 2d, Supplemental Fig. 6**). This indicated that Akita learned sequence features specifying an orientation-specific grammar of the CTCF sites most crucial for genome folding.

**Figure 2:**
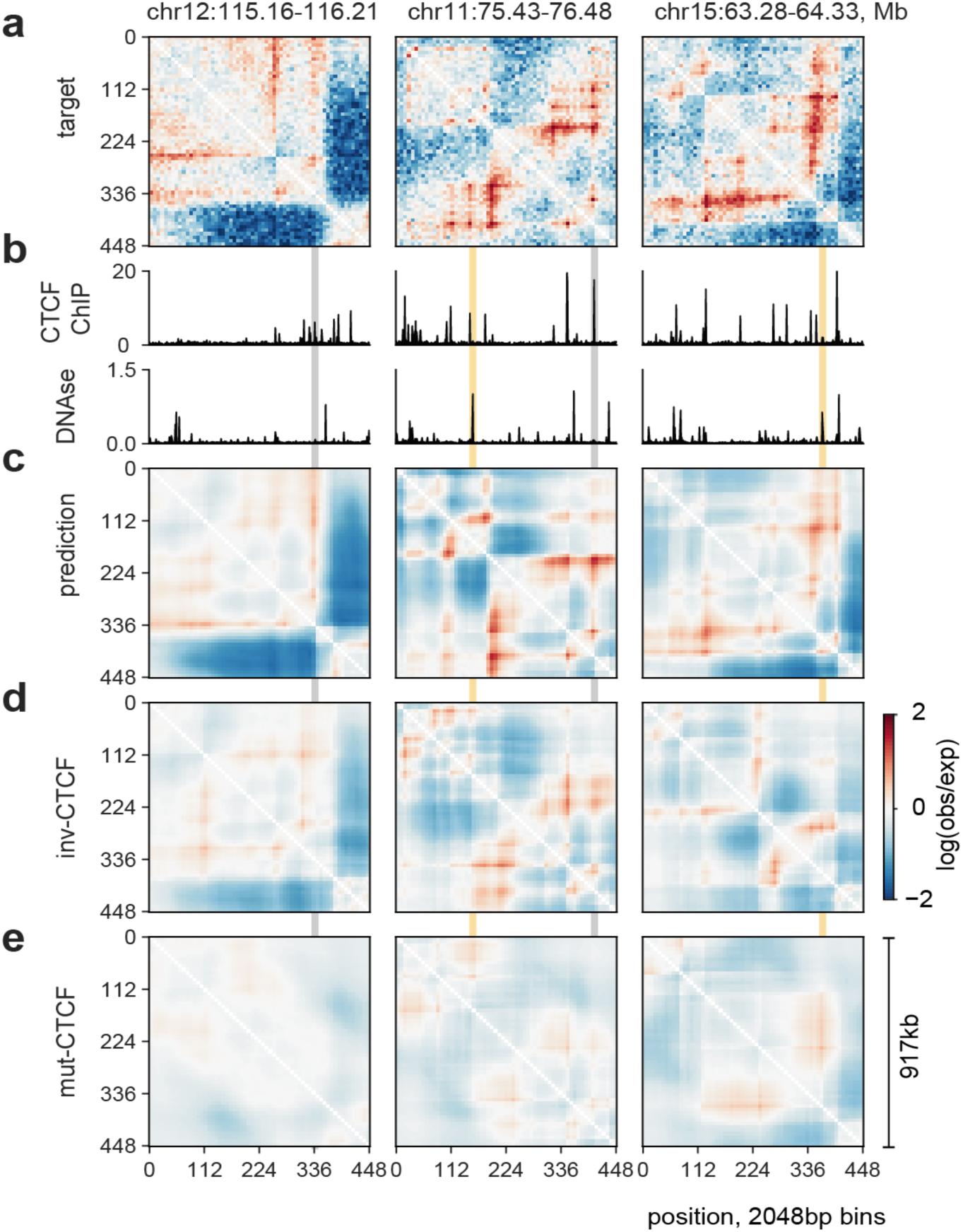
Akita learns a complex grammar for genome folding. **a**. Log(observed/expected) target HFF maps for three different genomic regions in the test set, binned to 2048bp. **b**. Binned profiles at 2048bp for CTCF ChIP-seq fold-change over control and DNAse density, data downloaded from the ENCODE data portal^31^. **c**. Predictions for the same three regions. **d**. Predictions for inverting all CTCF motifs in each region. Note that patterns are perturbed relative to (c), and have greater saliency of patterns as comped with (e). **e**. Predictions for random mutagenesis of all CTCF motifs within each region, averaged over ten instances. Grey shading shows regions with CTCF binding that are disrupted in these maps, and yellow shading shows regions with high DNAse but low levels of CTCF binding that are boundaries of residual structures after CTCF motif mutagenesis.

To explore the role of CTCF for Akita’s predictions genome-wide, we mutagenized the CTCF motifs in each region of the test set. The majority of mutagenized regions showed weaker locus-specific patterns (**Fig. 3a**), reminiscent of changes seen experimentally following acute CTCF degradation^21,22^. Performing a similar mutagenesis for each motif in the JASPAR transcription factor database^23^ revealed that CTCF had the strongest impact. The second largest effect was for CTCFL, which binds a very similar motif to CTCF but is typically inactive in somatic cells. For the remaining motifs, mutagenesis either imperceptibly disrupted genome folding or the predicted impact directly tracked the number of overlaps with CTCF motif positions (**Supplemental Fig. 5**). These results argue that no other transcription factor with a known motif plays as large of a role as CTCF for genome architecture, and that CTCF-independent aspects of genome architecture emerge from a combinatorial interplay between different DNA-binding factors.

We next investigated Akita’s ability to predict how genetically engineered mutations alter genome folding. As Akita makes predictions for 1Mb sequences and is not influenced by information beyond this window, we sought an example where a <100kb variant had a dramatic effect on genome folding. At the *Lmo2* locus in HEK293T cells^24^, two domains are separated by a boundary positioned at a cluster of three CTCF-bound sites (**Fig. 3C**). In cells with a ∼25kb deletion encompassing this boundary, the two domains merge. Making the same deletion *in silico* recapitulated this effect in the predicted Hi-C map (**Fig. 3C**). Leveraging Akita’s ability to rapidly assay sequence perturbations, we examined a combinatorial set of *in silico* deletions in the *Lmo2* locus (**Supplemental Fig. 7**). We found that deleting any individual CTCF site minimally altered predictions. Our model thus predicts this boundary is formed by redundant CTCF sites, a phenomenon observed experimentally in other genomic locations^3,4^.

**Figure 3:**
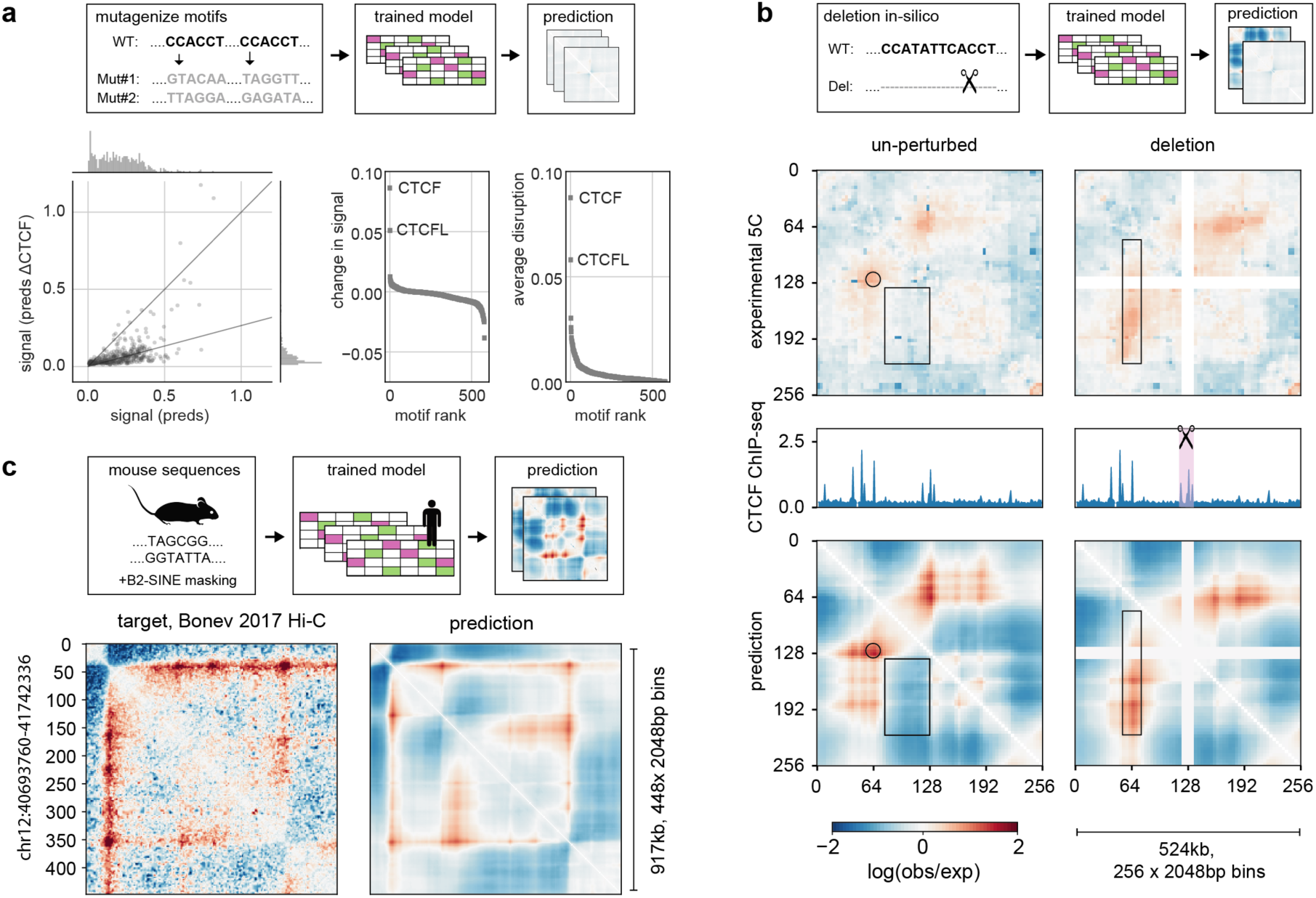
Applications of Akita. **a. *In-silico* motif mutagenesis**. *Left*: Predicted map signal strength before versus after mutagenizing all CTCF motifs, for each region in the test set for HFF model output. Map signal strength measured by<*pred* ^2^>. Akita predicts that mutagenizing CTCF motifs leads to more uniform maps, shown by the lower dynamic range after mutagenesis, 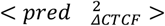 *Middle*: Change in map signal strength, measured by the difference of the mean squared values before versus after mutagenizing each motif in JASPAR^23^, 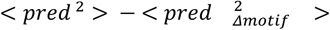. Positive values indicate lower signal after mutagenesis, as for CTCF. *Right*: Average disruption, measured by the mean-squared differences between predictions before versus after mutagenizing each motif in JASPAR, <(*pred* − *pred*) _*Δmotif*_)^2^ >. By this metric, CTCF mutagenesis is more than three times as impactful as mutagenesis of any other motif besides CTCFL. **b. Predicting a genetically engineered deletion**. *Top:* Experimental^24^ log(observed/expected) 5C data in HEK293T cells for WT (*left*) and a CRISPR/Cas9-mediated deletion of a ∼25kb boundary region (*right*) at the *Lmo2* locus for a 2^19^ bp region centered at the deleted boundary (chr11:33752474-34276762). In wild-type cells (*left*), this region displays a peak at the boundary (circle) between two ∼130kb domains that are relatively insulated from each other (rectangle), separated by a boundary that overlaps a cluster of three CTCF-bound sites. In cells where this boundary has been deleted (*right*), the two domains merge and display a flare of enriched contact frequency (thin rectangle). *Middle*: *CTCF* profiles for HEK293T^24^. *Bottom*: Computational predictions for WT (*left*) and deletion (*right*) of the boundary, showing similar changes. **c. Predicting mouse genome folding**. To further test Akita, which we trained on human datasets, we considered its accuracy for predicting mouse genome folding in mESCs. We found Akita often made overly pronounced predictions for mouse data, and this bias correlated with the number of SINE-B2 elements in the region. Plots show (*left*) experimental mESC Hi-C data^26^ and (*right*) computational prediction for one well-predicted region after mutagenizing SINE-B2 elements. See **Supplemental Fig. 8** for quantification across the mouse genome.

Given similar overall human and mouse genome folding^25^, we reasoned the mouse genome could provide evolutionarily perturbed sequences to further test Akita (**Fig. 3B**). Using mouse DNA sequences as input, we compared predictions from our human-trained model (hESC output) with mESC Hi-C data^26^. These cross-species predictions generally recapitulated mouse genome folding (**Supplemental Fig. 8**, median Spearman R: 0.50). Intriguingly, poorer predictions had more B2 SINE elements, which dramatically expanded in murid lineages and carry CTCF sites^27^. Mutagenizing B2 SINE elements improved our predictions for mouse genome folding (median Spearman R 0.55 vs 0.50). This suggests either the mouse genome specifically mitigates these elements, or Akita did not learn their true influence due to the lack of B2 SINEs in the human genome. These results are consistent with recent observations that the ChAHP complex hinders CTCF binding within murine B2-SINE elements^28^ and highlight opportunities for sequence-based modeling to uncover species-specific regulatory strategies.

An appealing hypothesis for future work is that neural networks with layers that better reflect the molecular and physical mechanisms organizing genomes will make more accurate and generalizable predictions. For the initial layers, convolutions naturally extend^11–13^ position weight matrix approaches for capturing the biophysics of protein-DNA interactions. The architectures and layers that might best reflect the process of loop extrusion, believed to organize mammalian interphase chromosomes,^29^ or other mechanisms of genome organization remain open questions. The near future promises exciting progress: recently, a similar CNN model, deepC, was posted to *bioRxiv*^*30*^. While deepC has a similar ‘trunk’ to Akita, it differs greatly in the architecture of the ‘head’, data pre-processing, and training schemes (**Supplemental Note 2**). Future work will benefit from comparing these approaches, continuing to explore the space of alternatives, and incorporating high quality data as it becomes available.

In summary, we present Akita, a model that predicts genome folding using only DNA sequence as an input. In the future, we envision that end-to-end sequence-to-genome-folding approaches will advance our ability to design functional screens, model enhancer-promoter interactions, prioritize causal variants in association studies, and predict the impacts of rare and *de novo* variants.

## Supplemental Figures

**Supplemental Figure 1:**
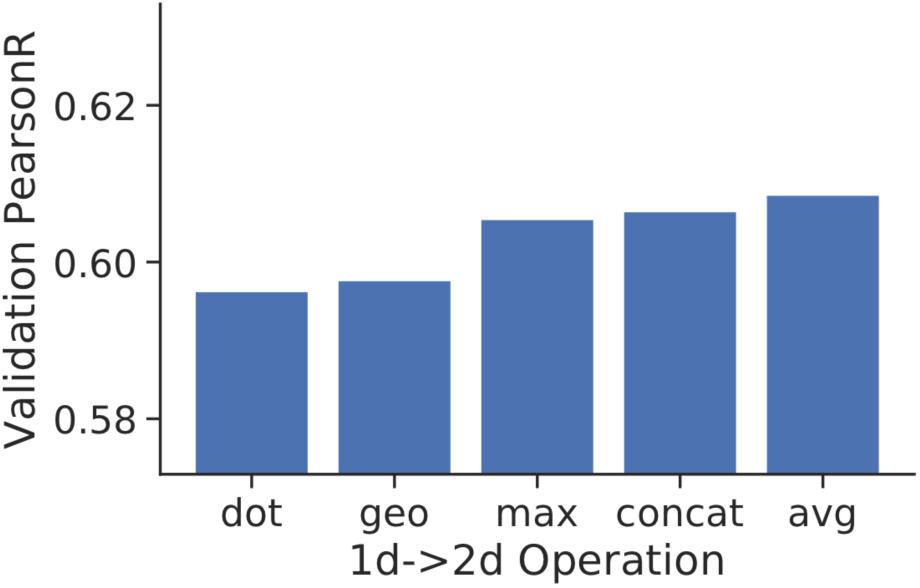
Transforming 1D representations to 2D via averaging produces better models than alternative operations. We considered the following operations to transform 1D vector representations derived from the DNA sequence to 2D for Hi-C prediction, holding all other hyper-parameters constant. For every pair of vectors *o*_*i*_ and *o*_*j*_ for 1D sequence positions i and j, we computed vector *t*(*i, j*)via: (1) “dot”: Element-wise multiplication between each vector position, *t*(*i, j, k*) = *o*_*i*_(*k*)*o*_*j*_(*k*). (2) “geo”: Addition of one to all vector values, element-wise multiplication between each position, square root of each position, subtraction of one from all vector values,

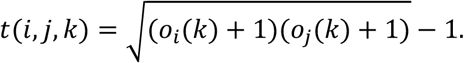 (3) “max”: Element-wise max between each vector position, *t*(*i, j, k*) = *max*(*o*_*i*_(*k*), *o*_*j*_(*k*)). (4) “concat”: Concatenate the two vectors, *t*(*i, j*) = [*o*_*i*_, *o*_*j*_]. (5) “avg”: Element-wise mean between each vector position, *t*(*i, j, k*) = (*o*_*i*_(*k*)+*o*_*j*_(*k*))/2.

**Supplemental Figure 2:**
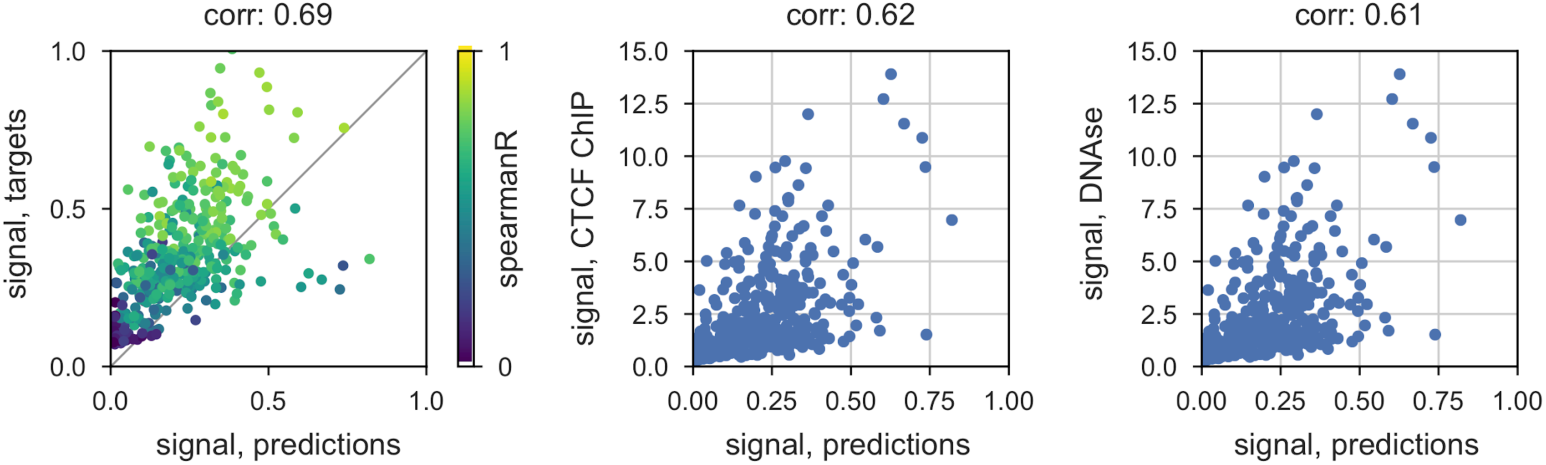
Correlations between the strength of Akita’s predictions, strength of experimental patterns, CTCF, and DNAse. *Left*: Map signal strength, measured by mean of squared map values, for predictions versus targets. In regions with more complex features, Akita tends to make more complex predictions. *Middle*: Signal strength for predictions versus signal strength for CTCF ChIP-seq, measured by mean squared profile values. Akita predicts more prominent locus-specific patterns in regions with greater CTCF binding. *Right:* Signal strength for predictions versus DNAse-seq.

**Supplemental Figure 3:**
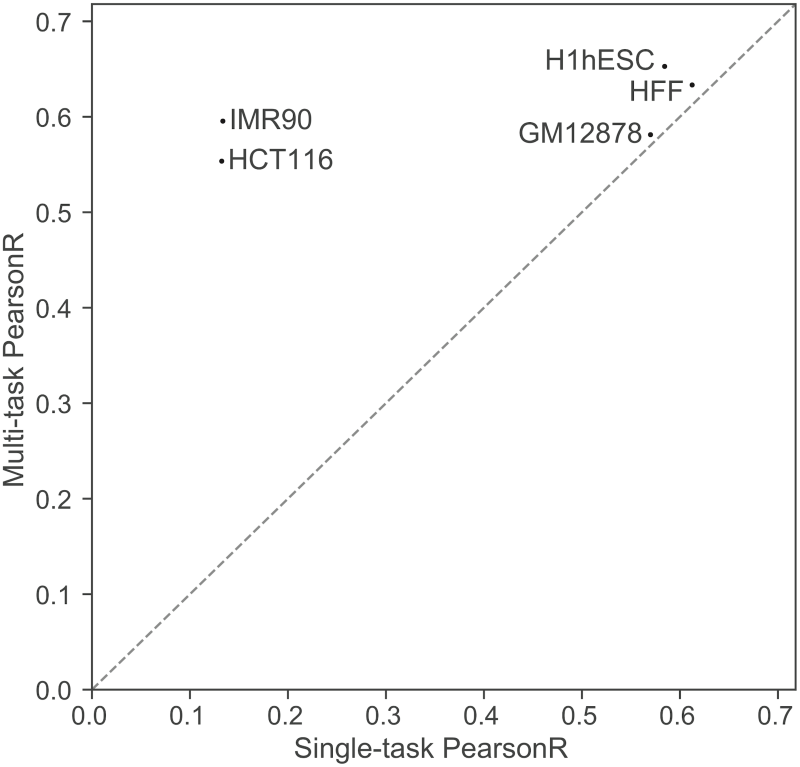
Multi-task training improves accuracy relative to single dataset training. We trained Akita models for each of the five datasets alone and compared PearsonR on the test set to the jointly trained multi-task model. Multi-task training greatly benefitted IMR90 and HCT116 learning, and also boosted accuracy for the other datasets.

**Supplemental Figure 4:**
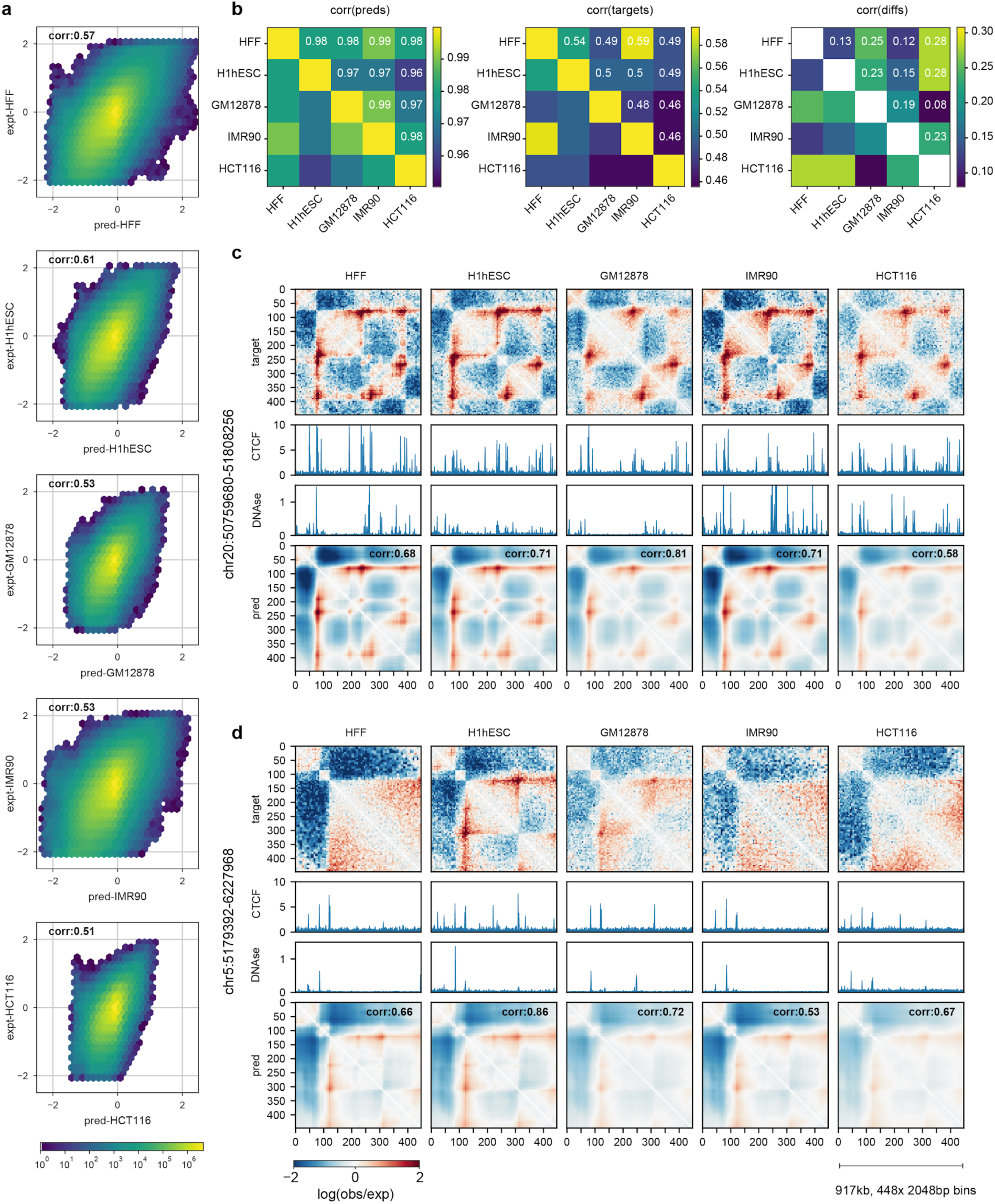
Model displays limited cell-type specificity in predictions. **a**. Predicted versus experimental log(observed/expected) values for each bin pair in every region of the test set, separately for each target. This shows predictions are correlated with experimental data across cell types. Color shows log10 number of bin pairs for each set of predicted versus experimental values. **b**. Considering every region in the test set across cell types, we find: ***Left*:** models make highly correlated predictions for different cell types. ***Middle***: genome folding assayed experimentally is correlated, but less so. ***Right***: predicted differences across cell types from our models correlate, albeit weakly, with observed differences. Note different scales for Spearman R. **c**. Example of a region showing largely consistent folding across cell types (chr20:50759680-51808256) for targets and predictions. Tracks show binned CTCF ChIP-seq fold-change over control, and DNAse-seq density. **d**. Example of a region showing gains and losses of specific features across cell types (chr5:5179392-6227968) at bin ∼300. While the predicted differences across cell types from models correlates with observed differences (**b, *right***), our predictions are not particularly visually distinct for different cell types (**c,d**). At present, our models appear to primarily tune the dynamic range for the entire prediction, rather than predicting gains and losses of a subset of features (**d**). Also note in **(d**) that CTCF is still bound in HCT116 in this region as determined by ChIP-seq, despite the loss of a strong boundary around bin 300. In the future, we hypothesize that a greater number of high-resolution genome folding datasets will enable our models to learn more cell-type specific representations of genome folding, as is currently possible for ATAC-seq and CAGE data ^16^.

**Supplemental Figure 5:**
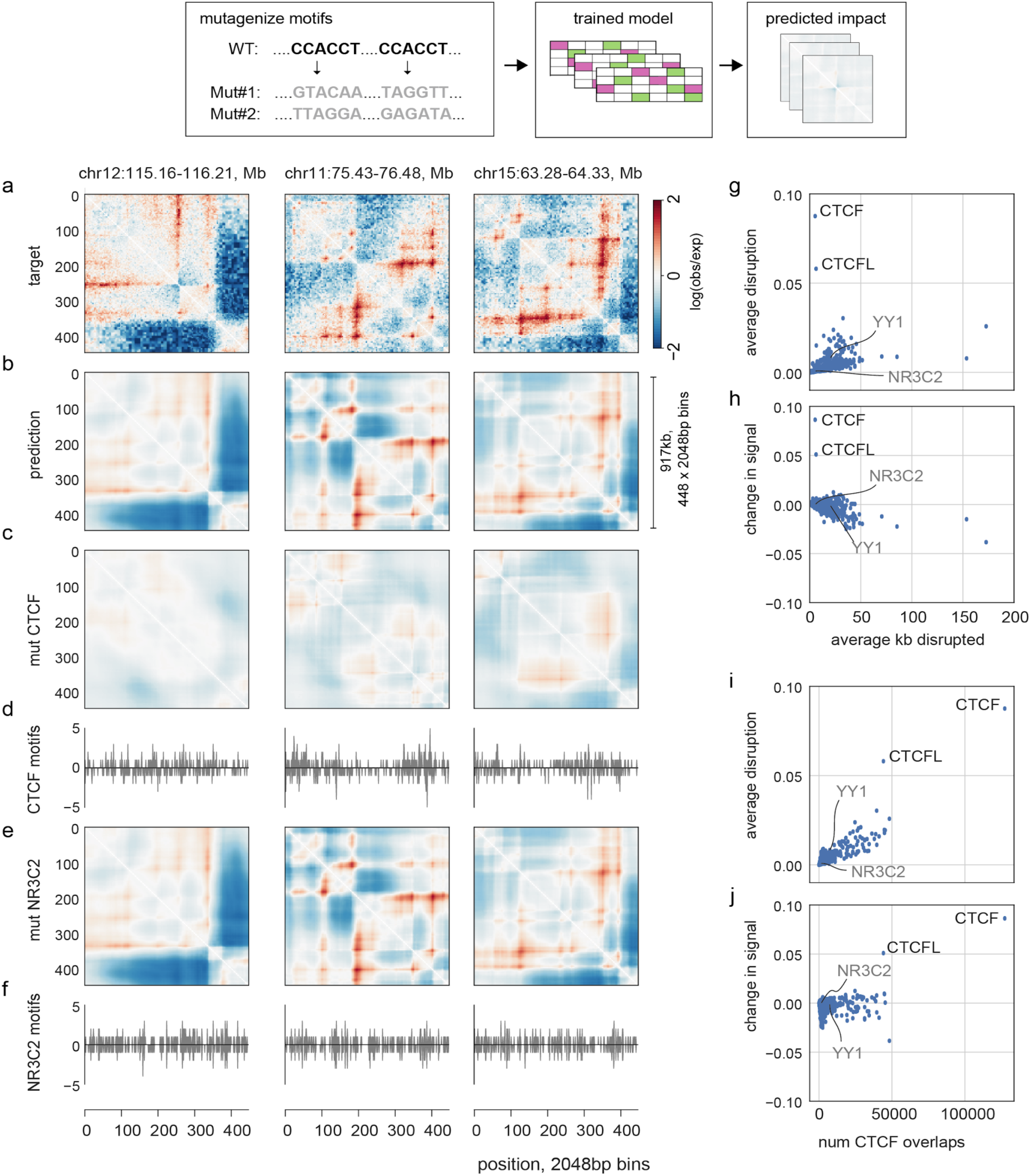
*In silico* mutagenesis enables rapid screening of transcription factor influence on genome folding. **a**. Experimental HFF Micro-C target data for three regions in our held-out test dataset. **b**. Predictions for these regions. **c**. Predictions for these regions after randomly mutagenizing all CTCF motifs in these regions, averaged over 10 random samples. **d**. Number of CTCF motifs per 2048bp bin. CTCF motif matches obtained from JASPAR ^23^, and profiles computed separately for the number of motifs on the positive strand (>0) and negative strand (<0). **e**. Predictions for these regions after randomly mutagenizing all NR3C2 motifs in these regions, averaged over 10 random samples. NR3C2 has a similar number of base pairs per region perturbed as CTCF, but little impact on Akita’s predictions. **f**. Positions of positively oriented and negatively oriented NR3C2 motifs. **g**. Average disruption, mean((pred-pred_mut_)^2^), versus the average number of kb perturbed per region. Note that YY1, suggested to be involved in genome folding^32,33^, is predicted to have little aggregate genome-wide impact following motif mutagenesis. This suggests YY1 may operate at a subset of loci in certain developmental contexts^32^, or its influence depends on the presence of nearby CTCF motifs. **h**. Change in signal, mean((pred)^2^) - mean((pred_mut_)^2^), versus the average number of kb perturbed per region. This reveals a trend towards negative scores for motifs with many occurences. **g**. Average disruption versus the total number of overlaps with CTCF motifs. This shows that the next highest scoring motifs after CTCFL likely have large predicted impacts due to overlapping CTCF binding sites, rather than independent effects. **h**. Change in signal versus the total number of overlaps with CTCF motifs.

**Supplemental Figure 6:**
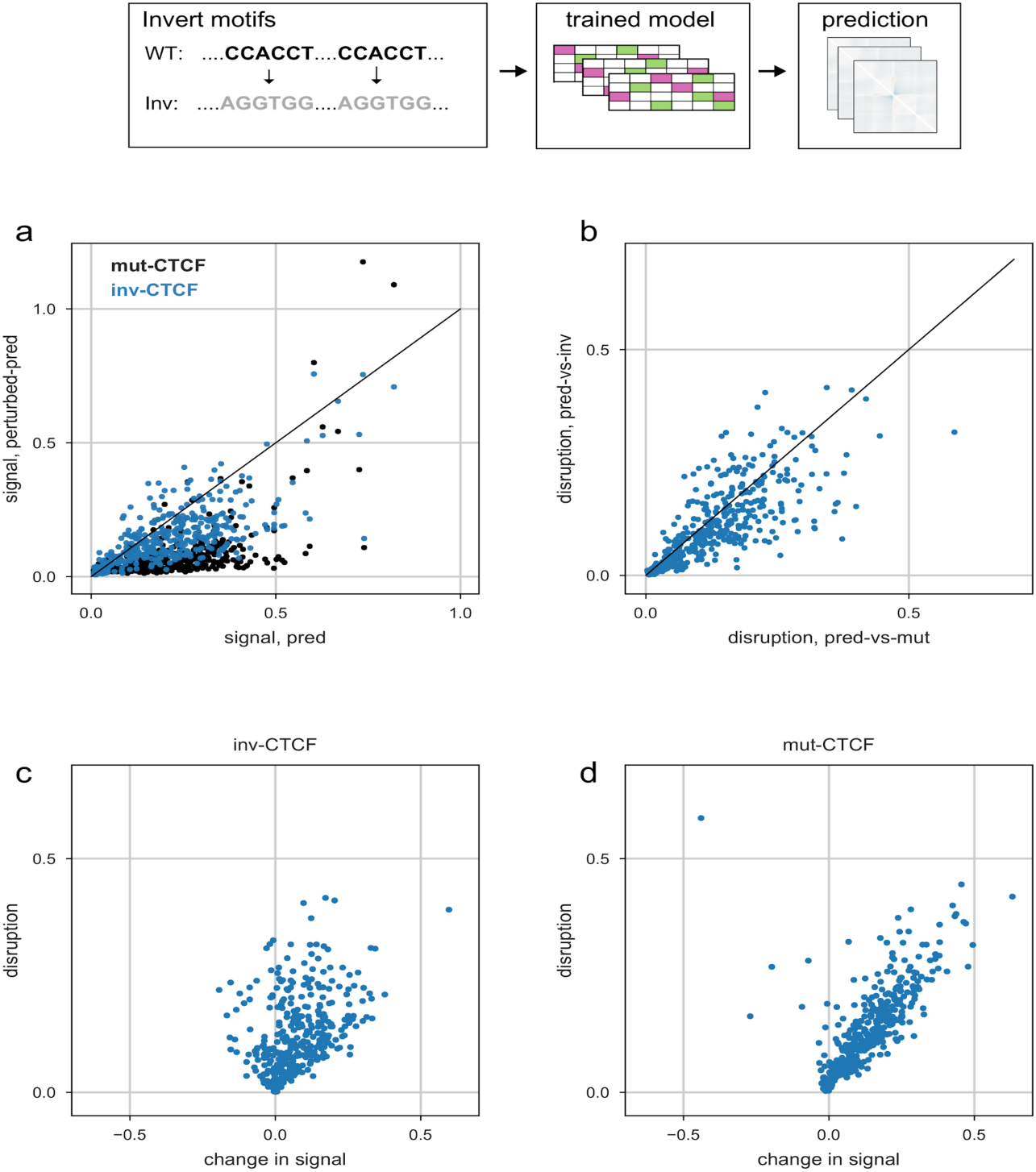
Akita learns an orientation-specific role for CTCF. **a**. Predicted map signal strength before versus after *in silico* perturbations, either for mutagenizing all CTCF motifs (black) or inverting all CTCF motifs (blue). Points show each region in the test set. Signal strength quantified by mean squared map values. Inversions show smaller perturbations to overall signal strength. **b**. Average disruption for mutagenizing all CTCF motifs or inverting all CTCF motifs. Inversion disruption maps to a similar extent as mutagenesis. Jointly with (a), Akita thus predicts changing motif orientation largely alters the positioning of contact patterns, rather than their overall salience across the genome. **c**. Change in signal strength versus disruption for inverting all CTCF motifs, mean((pred)^2^) - mean((pred_inv-CTCF_)^2^). Points show each region of the test set. This indicates that while motif inversions can greatly change the pattern of interactions (disruption), they can both increase (-) and decrease (+) the salience of contact map patterns. **d**. Change in signal strength versus disruption for mutagenizing all CTCF motifs in each region of the test set, mean((pred)^2^) - mean((pred_mut-CTCF_)^2^). Positive changes in signal strength upon mutagenesis shows these perturbations largely decrease pattern strengths in predicted maps.

**Supplemental Figure 7:**
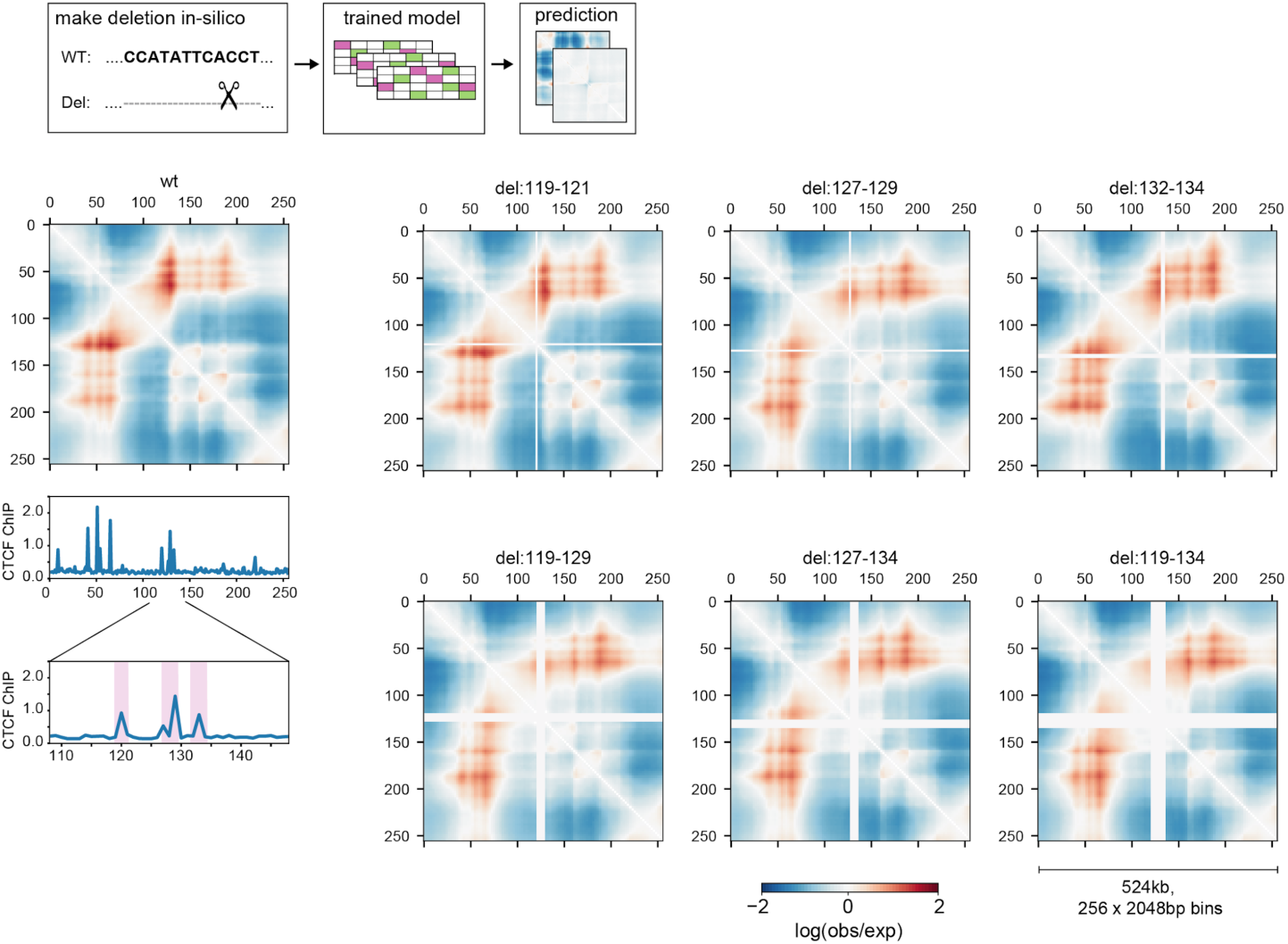
Model predicts a redundant boundary at Lmo2. *Left*: Predicted genome folding for unperturbed Lmo2 locus above the CTCF ChIP-seq profile for the region. Predictions in this figure used hg19 sequence as input and Akita’s output for HFF Micro-C ^20^. *Right*: Numbers above maps indicate the (start,end) position of bins that were deleted, highlighted by purple shading on the zoomed-in CTCF ChIP-seq profile below the predicted WT map. Akita predicts that deleting bins encompassing individual CTCF peaks (top row) would only mildly alter genome folding, and deletion of all three (bottom right) would be more impactful than either pair (bottom left and middle).

**Supplemental Figure 8:**
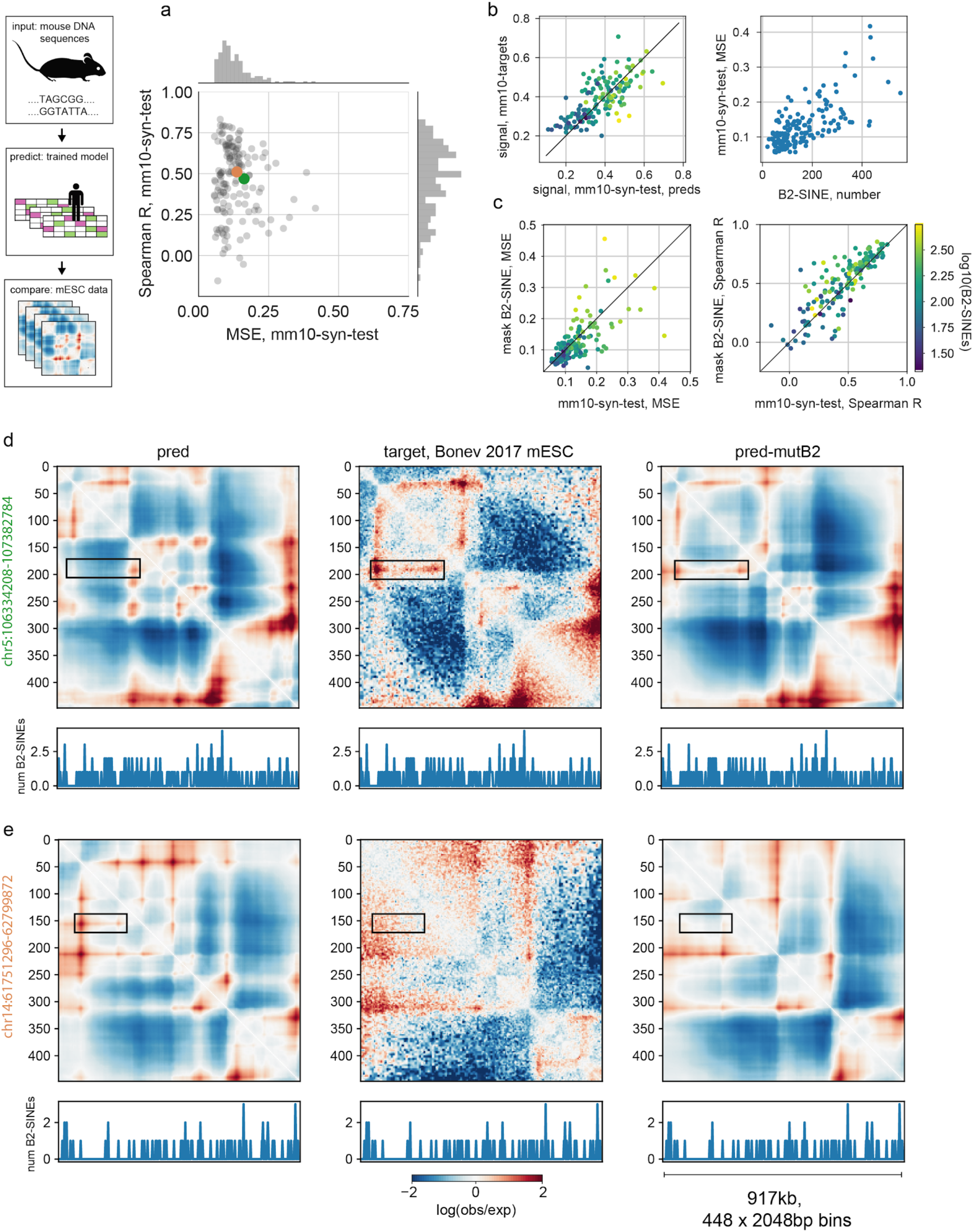
Cross-species predictions reveal impact of B2 SINE elements on genome folding in mouse embryonic stem cells. **a**. MSE-vs spearmanR for 156 mouse regions that overlap regions syntenic to the human test set (mm10-syn-test). The target Hi-C data was from mouse embryonic stem cells ^26^, mapped to mm10 and processed similarly to human datasets previously. Predictions in this figure were made using mm10 sequence as input and Akita’s output for the H1hESC Micro-C dataset ^20^. **b**. (*left*) Signal strength of predictions versus targets, for mm10-syn-test, calculated as the mean squared values in each map. The model trained on human data makes many overly strong predictions in mouse relative to the experimental data (see **Supplemental Fig. 2** for similar comparison with human experimental data). (*right*) Squared error between targets and predictions correlates with the number of B2-SINE elements in the region (from RepeatMasker^34^). **c**. Masking B2-SINE elements improved MSE for 93/156 predictions (*left*), and Spearman R for 106/156 predictions (*right*). This suggests the mouse genome has ways to mitigate the impact of its numerous B2-SINE elements on genome architecture. **d, e**. Examples of improved predictions for two regions from the mm10-syn-test set after masking B2-SINEs, with the total number of B2-SINE elements per bin in the region displayed below each map. Initial predictions indicated in (a) with orange and green dots. **d**. chr5:106334208-107382784 (deltaCorr:0.26, corrMutB2:0.72). Rectangle highlights a feature that is incorrectly predicted to be absent prior to masking B2-SINEs, and is correctly predicted following masking B2-SINEs. **e**. chr14:61751296-62799872 (deltaCorr:0.18, corrMutB2:0.69). Rectangle highlights a feature that is incorrectly predicted to be present prior to masking B2-SINEs, and is correctly predicted following masking B2-SINEs.

## Methods

### Code availability

All code used for training Akita available at: https://github.com/calico/basenji/tree/tf2_hic/. Trained Model at: https://github.com/calico/basenji/tree/tf2_hic/manuscripts/akita.

### Training Data

To obtain Hi-C data conducive for convolutional neural network learning, we reprocessed five of the highest-quality publicly available human Hi-C and Micro-C datasets to 2048bp (2^11^ bp) bins using *distiller* (https://github.com/mirnylab/distiller-nf)^35^ to map to hg38 and *cooler*^*36*^ to perform genome-wide iterative correction^37^.

| Target | Reference |
| --- | --- |
| HFF Micro-C | Krietstein et al., 2019<sup>20</sup> |
| H1hESC Micro-C | Krietstein et al., 2019<sup>20</sup> |
| GM12878 | Rao et al., 2014<sup>38</sup> |
| IMR90 | Rao et al., 2014<sup>38</sup> |
| HCT116 | Rao et al., 2017<sup>39</sup> |

To focus on locus-specific patterns and mitigate the impact of sparse sampling present in even the currently highest-resolution Hi-C maps, we: adaptively coarse-grain, normalize for the distance-dependent decrease in contact frequency, take a natural log, clip to (−2,2), linearly interpolate missing bins, and convolve with a small 2D gaussian filter (sigma=1, width=5). The first through third steps use *cooltools* functions (https://github.com/mirnylab/cooltools). Interpolation of low-coverage bins filtered out in typical Hi-C pipelines was crucial for learning with log(observed/expected) Hi-C targets, greatly outperforming replacing these bins with zeros.

To prepare the Hi-C data for training, we divided the human genome into large virtual contigs and assigned them to training, validation, and test sets with an 80/10/10 split. We broke the chromosomes at assembly gaps, large unmappable regions, and consecutive stretches of ≥10 filtered-out Hi-C bins (in any target dataset). Within the contigs, we extracted 2^20^ bp (∼1Mb) sequences, striding by 2^18^ bp (∼262kb) for the training set and 2^19^ bp (∼524kb) for the validation and test sets. This procedure resulted in 7,008 training, 419 validation, and 413 test sequences.

### Model architecture

We created a neural network architecture to predict 2D Hi-C maps from 1D DNA sequences that consists of two major components. First, we process the 1D DNA sequence using a ‘trunk’ that applies a series of convolutions, following previous work on convolutional neural networks for DNA sequence analysis. Second, we applied a ‘head’ that transforms the 1D representations to 2D for Hi-C prediction. We implemented the model using the Basenji software^16,17^, which is written in Tensorflow^40^ and Keras^41^.

More specifically, the ‘trunk’ includes:

1. Convolution with 96 filters of size 11-by-4 to transform the 1-hot encoded DNA sequence followed by batch normalization, ReLU, and width 2 max pooling.
2. Convolution tower that iteratively performs convolution with 96 filters of width 5, batch normalization, ReLU, and width 2 max pooling to arrive at 512 vector representations of the sequence in 2048bp windows.
3. Dilated residual convolution tower that iteratively performs dilated convolution with geometrically increasing dilation rate, adding the new representation back into the old. This block spreads information about relevant sequence elements and global context across the sequence^16^.
4. Bottleneck width 1 convolution with 48 filters.

To convert these 1D representations to 2D for the Hi-C ‘head’, we averaged the representations for every pair of genomic bins *i* and *j*. This operation transforms a tensor with dimensions [512 length, 48 filters] to a tensor with dimensions [512 length, 512 length, 48 filters]. We also concatenated a positional encoding of the distance between bins, abs|i-j| and applied a (1,1) convolution block to finalize the transition to 2D. Next, we treat this map as a 2D image and run multiple layers of dilated residual 2D convolutions with geometrically increasing dilation rate, re-symmetrizing after each step. Finally, we apply one last linear transformation to make predictions for the 5 datasets.

Intuitively, the initial transformation to 1D to 2D should be able to recognize genomic features with important relationships for Hi-C prediction, such as two boundary elements, and the subsequent 2D convolutions serve to disseminate that recognition to the surrounding region. Intriguingly, similar sequence-to-map architectures have recently been successful for protein contact map prediction^42^.

### Training Approach

We computed a mean squared error loss from the targets and predictions, considering only the upper triangular portion of the matrixes. We fit the model parameters using stochastic gradient descent with momentum for ∼60 epochs, taking steps in batches of 2 sequences.

Data augmentation was critical to avoid overfitting and maximize generalization accuracy to unseen sequences. Each time that we processes a sequence, we stochastically shifted input sequences by up to +/-11 bp and reverse complemented the DNA and flipped the Hi-C map.

We stopped training when validation loss had not improved for 12 epochs, and we took the model parameters that had achieved that minimum validation loss forward as the final model. We performed a search over learning rate, momentum, gradient norm clipping, dropout probability, and convolution filters using the Dragonfly Bayesian optimization toolkit [https://github.com/dragonfly/dragonfly]^43^.

### Comparison with 1D features

For comparison to 1D features of the epigenome, we downloaded processed bigWigs for the relevant cell types from the ENCODE data portal ^31^ and binned them into 2048bp profiles.

### *In silico* motif mutagenesis

To perform *in silico* motif mutagenesis, we intersect our test set regions with motif positions using bedtools ^44^. We then generate multiple randomized sequences, where DNA sequence at positions of motifs with randomly generated DNA sequences of the same length. We then calculate the *average disruption* as mean((pred - pred_Δmotif_)^2^), and the *change in signal* as mean(pred^2^) - mean(pred^2^ _Δmotif_). Motifs names were plotted with adjustText (https://github.com/Phlya/adjustText)^45^. Maps in **Fig. 2** shown as averages over 10 randomized sequences, JASPAR-wide analyses in **Fig. 3a** averaged over 3 randomized sequences.

### *In silico* CTCF motif inversions

We perform in silico motif inversions similarly to motif mutagenesis for determining intersections. We then merge overlapping motifs and replace sequences in these intervals with their reverse complements.

### Predictions for mouse DNA sequences

To test the accuracy of Akita’s predictions for mouse DNA sequences, we obtained mESC Hi-C data from Bonev et al., 2017 ^26^, mapped reads to mm10, and otherwise processed the data as for human datasets. Positions of B2-SINE elements were downloaded from UCSC (from RepeatMasker^34^). B2-SINE mutagenesis was performed as described for motifs.

### 5C data processing

To test Akita’s ability to predict experimentally induced deletions, we obtained processed 5C data for the *Lmo2* locus from Hnisz et al., 2016^24^, re-binned fragments to 2048bp bins, and otherwise performed the same processing into log(observed/expected) maps as for Hi-C data above.

### *In silico* deletions

As Akita makes predictions for fixed input size, to make a deletion *in silico* we must both remove the DNA sequence we hope to delete and supply the model with an equal amount of additional DNA sequence. Here we centered on the position of the deletion and symmetrically extended the start and end to maintain the size of the input.

## Supplemental Notes

### Supplemental Note 1: lessons from previous architectures

Before arriving at the model described above, we considered various alternative designs in the space of possible model architectures and data pre-processing schemes.

Some of our early attempts at predicting 3D genome folding from sequence involved predicting slices of the Hi-C matrix (‘virtual-4C’) directly from the outputs of the trunk, with the idea that predicting a contact vector of length N could require fewer parameters in the ‘head’ than predicting a contact map of size NxN. While such virtual-4C models readily learned boundaries, we found they failed to learn sharp peaks. We also found that predictions of these models were often asymmetric (i.e. pred_i,j_ != pred_j,i_), likely because the virtual-4C architecture we considered did not encode a symmetry constraint.

We also tested the performance of models that replace the dilated convolution layers in the trunk with bidirectional LSTMs, popular layers for capturing long-range dependencies in natural language processing, while preserving roughly the same time per training epoch. This architecture performed slightly more poorly on both training and validation data, and we did not pursue it further. We also explored the utility of separable convolutions to reduce the number of parameters in the convolutional tower: we found little benefit in the rate of learning and a slight loss in accuracy.

### Supplemental Note 2: differences with DeepC

In Schwessinger et al.^30^, the authors report successful predictions of Hi-C maps at 10kb resolution using a similar deep convolutional neural network approach, deepC. While deepC has a similar ‘trunk’ to Akita, it differs greatly in the architecture of the ‘head’, data pre-processing, and training schemes. First, deepC uses non-linearly quantile normalized targets, rather than log(observed/expected) targets, which may emphasize learning peaks rather than insulation. Second, deepC predicts a ‘zig-zag pole’ target directly with a dense layer from the output of their model’s trunk, which implicitly encodes distance but requires a large number of parameters, rather than predicting a dense patch of a map. Third, we focused on higher-resolution predictions (2048bp bins vs. 10kb bins). Finally, deepC currently requires pre-training a model on a large set of epigenomic profiles, and transferring weights to their full model. It is possible that strict transfer learning could limit the richness of representations that a deep CNN can learn for 3D genome folding; for example, a CTCF profile may not contain information about the directionality of motifs under its peaks, which is important for predicting genome folding.

## Acknowledgements

The authors thank Vikram Agarwal, Han Yuan, and Elphege Nora for detailed feedback. GF and KSP were funded by Gladstone Institutes, the National Heart, Lung and Blood Institute (grant #HL098179), and the National Institute of Mental Health (grant #MH109907).

